# Combination adjuvants drive long lived plastic Th17 cells that convert to multi-functional Th1 cells and protect mice against fungal infection

**DOI:** 10.64898/2026.04.03.716375

**Authors:** Marcel Wüthrich, Uju Joy Okaa, Cleison Ledesma Taira, Lucas dos Santos Dias, Bruce S. Klein

## Abstract

Th1 cells are viewed as a cornerstone of immunity to fungi and other intracellular pathogens. Despite the widely accepted role of Th1 cells in antifungal resistance, the development of protective strategies harnessing them is stunted by a limited understanding of how best to promote their development. We and others have reported a requisite role for Th17 cells in resistance to fungi. We have long been puzzled about how to reconcile seminal roles for both Th1 and Th17 subsets. Here we report that Th17 cells convert into polyfunctional Th1 cells producing multiple cytokines, including IFN-γ, TNF and GM-CSF when we used adjuvant formulations that include glucopyranosyl lipid adjuvant (GLA) to enhance antifungal immunity. GLA induced plastic Th17 cells that convert into polyfunctional Th1 memory cells.

## INTRODUCTION

Despite the successes in combating deadly bacterial and viral diseases (1) and the growing clinical need, there are currently no fungal vaccines licensed for human use (2). A major limitation for developing anti-fungal vaccines is the lack of highly protective antigens and adjuvants that promote long lasting protective immunity that relies on antigen-specific Th1 cells. Hence, here we investigated how optimal Th1 immunity develops given the importance and centrality of IFN-γ and type 1 cytokine producing CD4^+^ T cells in orchestrating antifungal defense.

We recently discovered *Blastomyces*-Eng2 (*Bl-*Eng2), a fungal glycoprotein in *Blastomyces dermatitidis (Bd)* that harbors potent antigenic (3) and adjuvant functions (4). Vaccination with *Bl-*Eng2 elicits cellular immunity and protects mice against experimental infection with this fungus. Analysis with a peptide-MHC tetramer specific to an immunodominant region of the antigen has demonstrated that vaccination and infection expands and recruits several hundred thousand T cells to the lung after infection (3). This subunit vaccine incorporates a conserved antigen and is protective in mice when given subcutaneously (SC), but not intranasally (IN)(3, 5). By using single cell transcriptome analysis, we observed that vaccine-induced protection was tightly associated with IFN-γ production whereas nonprotective vaccination revealed Th17 skewed responses (6). Aside from the largest populations of polarized, conventional Th1 and Th17 cell signatures these clusters were adjoined by a spectrum of 5 additional populations also expressing Th1- or Th17-related genes, or both (6). This fluidity of the dominant cytokine phenotypes in Th1/Th17 cells complicated assignments of strict Th archetypes and may align with evolving notions of Th cell cytokine plasticity, for instance in Th17 cells that convert their cytokine phenotype described elsewhere (7).

Based on our previous findings and study (6), we hypothesized that vaccination with the Eng2 subunit vaccine elicits plastic Th17 cells that convert to protective Th1 memory T cells. We formulated the GCP-*Bl*-Eng2 vaccine, which contains two adjuvants (GCP and Eng2), together with the TLR4 ligand Glucopyranosyl lipid adjuvant (GLA) that promotes differentiation of primed T cells towards the Th17 cell lineage and followed the development of memory T cells (8).

Herein, we report that 1) formulation of *Bl*-Eng2 with GCP and GLA engenders durable immunity and protects vaccinated mice against experimental infection long term for at least one year. 2) The addition of GLA to other adjuvants significantly augments the survival of vaccinated mice that have rested for one year before challenge. 3) The addition of GLA induces plastic Th17 cells that convert to multi-functional Th1 cells that produce TNF, GM-CSF and IFN-γ over the course of the T cell contraction and memory phases. 4) Adoptive transfer of plastic memory Th17 cells protects naïve mice against lethal fungal infection. 5) The potency of highly protective vaccine hinges not only on IFN-γ, but on multiple cytokines, including IFN-γ, TNF and GM-CSF underscoring the conclusion that polyfunctional Th1 cells confer high level protection.

In summary, we conclude that the combination adjuvants described here induce the generation of plastic Th17 cells that convert into multi-functional Th1 cells and augment memory and protective immunity to fungi.

## METHODS

### Sex as a biological variable

Both male and female C57BL/6 mice were used in vaccination experiments and similar outcomes were found.

### Fungi

Wild-type *Bd* strain ATCC 26199 was used for this study. *Bd* 26199 was grown as yeast on Middlebrook 7H10 agar with oleic acid-albumin complex (Sigma) at 39°C.

### Mouse strains

Male and female mice were 7 to 8 weeks old at the time of these experiments. Inbred wild-type C57BL/6 mice (stock #000664) obtained from Jackson Laboratories were bred at our facility. Male and female mice were 7 to 8 weeks old at the time of these experiments. A breeding colony of B6.129(SJL)-*Il17a^tm1.1(icre)Stck^*/J (stock #035717) knock-in/knock-out was bred to *Gt(ROSA)26Sor^tm1(EYFP)Cos^*(stock # 006148) reporter mice to allow fluorescent labeling of cells expressing IL17A. Resulting heterozygous IL-17a^cre/+^ /Gt(Rosa)26Sor^eYFP/+^ mice that had one functional copy of IL-17 and one copy of IL-17-Cre x flox for fate mapping by eYFP fluorescence were used in this study. Mice were housed and cared for in a specific-pathogen-free environment in our animal facility as per guidelines of the University of Wisconsin Animal Care Committee, who approved all aspects of this work.

### Generation and purification of recombinant *Bl*-Eng2

*Bl*-Eng2 was cloned and expressed in *Pichia pastoris* using standard recombinant techniques and have been described (9). Recombinant proteins were purified using Ni-NTA agarose (Qiagen) according to the manufacturer’s protocol and dialyzed against PBS. Quantity and purity of recombinant *Bl*-Eng2 was assessed by Bradford assay, SDS-PAGE and silver staining.

### Vaccination and fungal infection

Ten micrograms of *Bl-Eng2* were loaded into GCPs(10). In a few experiments 10 μl of Advax3 adjuvant (11) was combined with 10 μg of soluble *Bl*-Eng2 protein and adjusted to a final volume of 200 μl with phosphate-buffered saline (PBS). Monophosporyl Lipid A (GLA) was purchased from Avanti Polar Lipids, LLC and resuspended in DMSO at 3mg/ml. GLA was mixed with GCP-*Bl*-Eng2 at 5 μg/mouse. Mice were vaccinated subcutaneously (SC) with *Bl-*Eng2 full length protein three times, two weeks apart as described (12) with antigen loaded into glucan chitin particles (GCPs); controls in these experiments were GCPs loaded with mouse serum albumin (MSA) (Dr. Gary Ostroff, University of Massachusetts Chan Medical School). Two weeks after the vaccine boost, mice were challenged intratracheally (i.t.) with 2×10^4^ wild type *Bd* yeast in 30μl PBS. At day 4 post-infection, lung T cell responses were analyzed and lung CFU counted. Two weeks post infection, when control or unvaccinated mice were moribund, fungal burden was determined by plating lung CFU.

### Generation of MHCII tetramer

Tetramers for detection of *Bl*-Eng2-specific CD4^+^ T cells in C57BL6 were generated at the NIH Tetramer core facility at the Emory University in Atlanta, GA.

### ELISA

Cytokine concentrations in cell culture supernatants were determined by IFN-γ and IL-17A Duoset ELISA kits (R&D Systems, Minneapolis, MN) according to manufacturer’s instruction.

### CD4 T cell enrichment before sorting

Miltenyi LS columns on a quadroMACS magnet were used to enrich CD4^+^ cells from secondary lymphoid organs (SLO). Spleen and draining lymph nodes were harvested and mashed through 40 μm filters. Red blood cells were lysed with ACK buffer. Samples were washed with RPMI and resuspended in cold sorter buffer (PBS with 2% FBS) to a volume twice the size of the pellet. Two microliters of Fc block was added to each sample and incubated for 5 min before adding anti-CD4 mAb coated microbeads. CD4 staining was done for 30 min at 4°C in the dark. After wash, samples were resuspendedin 3 mL of sorter buffer, filtered in 40 um filter, and added in pre-wet LS column. Columns were washed with cold sorter buffer thrice before eluting bound fractions. Fractions were stained with Invitrogen’s LIVE/ DEAD^TM^ stain and surface markers.

### Sorting of eYFP^+^ T cells

Following fluorescent labeling, cells from 10-15 animals from each vaccine group were combined into one tube each for cell sorting. eYFP^+^ cells were sorted into microcentrifuge tubes containing RPMI media on a FACs Aria using a 130 micron nozzle. The sorted cells (Live, Dump-, CD90.2^+^CD4^+^CD44^+^eYFP^+^) were collected directly into 1.5 ml microtubes. 72,000 eYFP^+^ T cells were adoptively transferred into naïve recipient mice.

### T-cell stimulation, and flow cytometry

Lungs were dissociated in Miltenyi MACS tubes and digested with collagenase (1 mg/mL) (Sigma) and DNase (1 μg/mL) (Sigma) for 25 min at 37°C. Digested lungs were resuspended in 5 mL of 40% percoll; 3mL of 66% percoll was underlaid (GE healthcare, cat# 17–0891–01). Samples were spun for 20 min at 2,000 rpm at room temperature. Lymphocytes in the buffy coat were harvested and resuspended in RPMI (10% FBS, 1% penicillin and streptomycin). For T-cell stimulation *ex vivo*, cells were incubated at 37°C for 5h with 5 μM peptide and 1μg anti-mouse CD28 (BD #553294). After 1h, BD GolgiStopTM (BD, cat# 554724) was added to samples. FACS samples were stained with Invitrogen’s LIVE/DEAD^TM^ stain and Fc block for 10 min at room temperature. Cells were stained with tetramer for 1h at room temperature, or for surface antigens (anti-CD45-AF488 (Biolegend: clone 30-F11 Cat #103122); anti-CD8-PerCpCy5.5 (Biolegend: clone 53.67 Cat #100734); anti-CD90.2-BV421 (Biolegend: clone 30H12 Cat #105341); anti-CD44-BV786 (Biolegend: clone IM7 Cat #103059); anti-CD4-BUV737 (BD Bioscience: clone RM4-5 Cat #612844) and dump channel antibodies (anti-CD11c-APC (Biolegend: clone N418 Cat #117310); anti-CD11b-APC (Biolegend: clone M1/70 Cat #101212); anti-NK1.1-APC (Biolegend: clone P136 Cat # 108710); anti-B220-APC (Biolegend: clone RA3-6B2 Cat #103212)) or intracellular targets (anti-IFN-γ-PE-Cy7 (Biolegend: clone XMG1.2 Cat #505826); anti-IL-17A-PE (Biolegend: clone TC11-18H10.1 Cat #506904); anti-IL-5-BV421 (Biolegend: clone TRFK5 Cat #504311); anti-IL13-eFlour450 (eBioscience: eBio13A Cat #48-7133-82) for 20 min at 4°C. Transcription factors were stained on primed, resting cells using the Foxp3 Transcription Factor Staining kit (ebioscience cat# 00–5523–00). All panels included a dump channel (Dump markers:CD11b, CD11c, NK1.1, CD19 B220). 50 μl AccuCheck Counting Beads (Invitrogen PCB100) were added to samples to determine absolute cell counts. Samples were acquired on a LSR Fortessa.

### Cytokine neutralizations

Mice were vaccinated with GCP-*Bl-*Eng2+GLA thrice and rested for 9 months post-vaccination. At the time of challenge and every other day thereafter, mice were treated intravenously with 250 μg of anti-IL-17A (BioXcell, clone 17F3, Cat# BE0173), anti-IFN-γ (BioXcell, clone XMG1.2, cat# BE0055), anti-TNF (BioXcell, clone XT3.11, cat# BE0058) and anti-GM-CSF (BioXcell, clone MP1-22E9, Cat# BE0259) or control rat IgG (BioXcell, polyclonal, Cat# BE0094).

### Statistical analyses

A 2-tailed, Mann-Whitney U test was primarily used to analyze the differences between 2 treatment groups for cytokine ELISAs, cytokine concentrations, calculations of the number of lung- infiltrating immune cells, and percentages of specific cytokine-producing T cells. In some instances, 1-way ANOVA was used when comparing multiple groups, and when a result was significant, a Tukey’s or Dunnett’s post hoc test was used to adjust for multiple comparisons. Differences in fungal burden (expressed as CFU) between 2 groups were analyzed using the Mann-Whitney U test for ranking data. For comparison of fungal burden among 3 or more groups of mice, the Kruskal-Wallis test, a nonparametric ranking method was used. Survival data were examined by the Kaplan-Meier test using log-rank analysis to compare survival plots as reported previously (6). A P value of 0.05 or less was considered statistically significant. Comparisons in many experiments yielded P values at or below the value of 0.05, however we consistently used only 1 asterisk throughout to denote any statistically significant difference, regardless of the exact P value below 0.05.

### Study approvals

Animal studies were conducted with adherence to protocol M005891 approved by the Institutional Animal Care and Use Committee of UW-Madison (IACUC). The studies were compliant with provisions established by the Animal Welfare Act and the Public Health Services Policy on the Humane Care and Use of Laboratory Animals.

## RESULTS

### Vaccination with *Bl*-Eng2 induces durable immunity against infection with *Bd*

Adjuvants function as antigen delivery systems and/or as immunopotentiators and modulators. Here, we formulated *Bl-Eng2* inside glucan-chitin particles (GCP) and studied the evolution of the ensuing antigen (Ag)-specific CD4^+^ T cell response and vaccine-induced resistance against *Bd*. At serial time points post-vaccination, we determined the number of tetramer-positive (**Fig. 1A+B**), and cytokine-producing (**Fig. 1A+C**) T cells in the circulating blood, the lung, the skin draining lymph nodes (dLN) and the spleen. The numbers of tetramer-positive and cytokine-producing T cells peaked at 14 days, contracted over the course of 100 days, and remained elevated for up to 400 days post-vaccination compared to unvaccinated mice. At 12 months post-vaccine, *Bd*- challenged mice had significantly fewer lung CFU compared to unvaccinated control mice (**Fig. 1D**). The reduction in lung CFU coincided with elevated numbers of tetramer-positive T cells (**Fig. 1E**) that produced IFN-γ and IL-17 (**Fig. 1F**). In summary, T cell memory specific for *Bl-Eng2* was durable and functional as measured by cytokine production and the ability to reduce lung CFU.

**Fig. 1:**
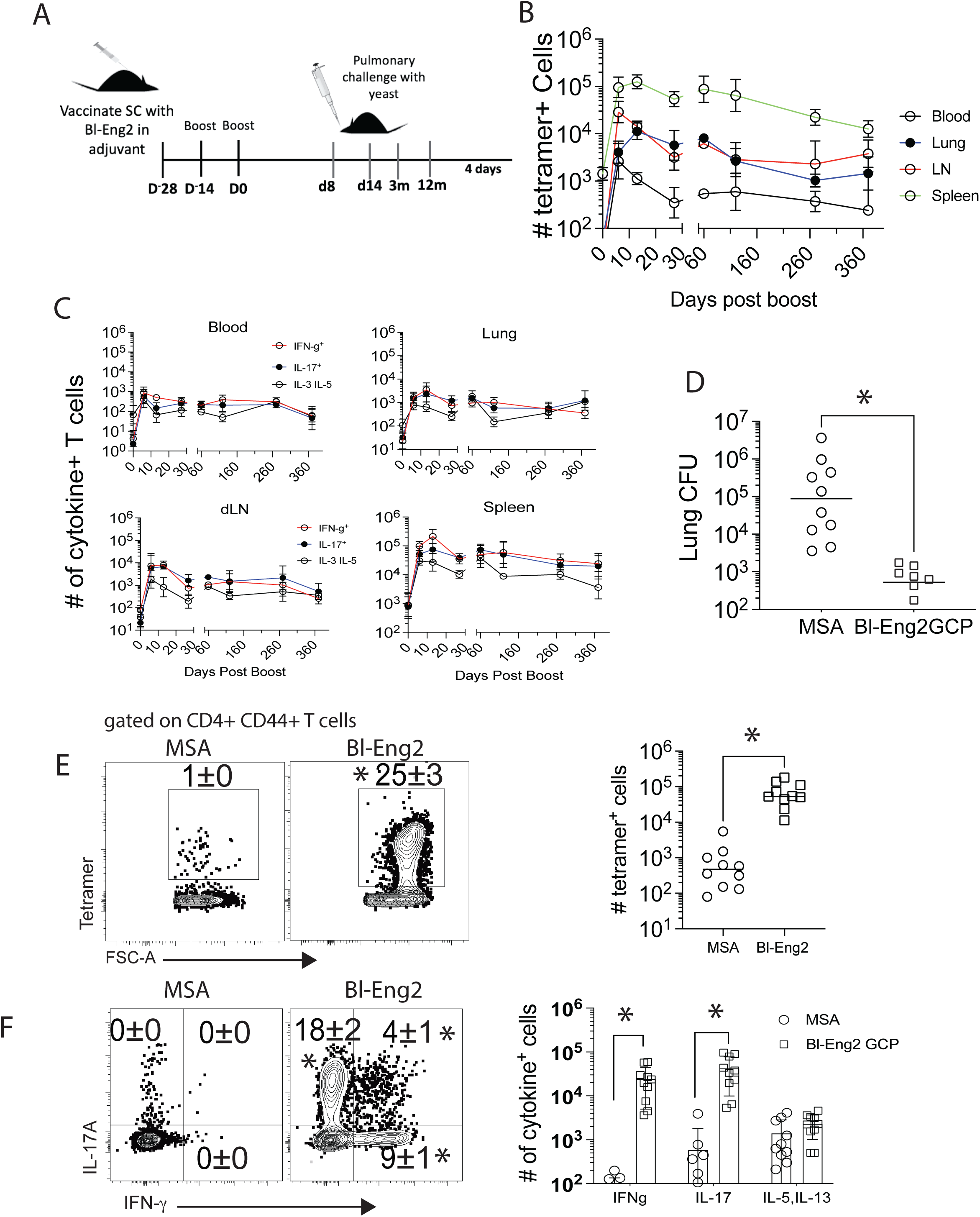
Vaccination with GCP-*Bl*-Eng2 engendered long lasting memory and protection against infection with *Bd* over 12 months. C57BL6 mice were vaccinated three times spaced two weeks apart and cellular immune responses monitored in the lung, skin draining lymph nodes, spleen and blood at serial time points post-vaccination (**A**). The number of tetramer positive T cells (**B**) and cytokine producing T cells (**C**) were enumerated for up to one-year post-vaccination. At twelve months post-vaccination, lung CFU (**D**), tetramer positive (**E**) and IFN-γ and IL-17 (**F**) producing T cells were analyzed. Dot plots are a concatenation of 5 mice/group, representing 2 independent biological experiments. The frequencies of cells are calculated as geometric means ± geometric SD. CFU from at least 10 mice/group is expressed as Log_10_ plotted with geometric mean ± geometric SD. *p<0.05, 2-tailed Mann-Whitney T test.

### Adjuvant combination augmented cellular immunity and vaccine induced protection

Engagement of multiple innate immune pathways may be vital in programming a potent and protective immune response (13, 14). We sought to test whether adjuvant combinations could result in improved cellular and vaccine immunity. We vaccinated mice with the *Bl-*Eng2 antigen and one of three combination adjuvants (**Fig. 2A**): 1) Advax3 containing inulin particles and the TLR9 agonist CpG driving a Th1 response (11), 2) Glucan-chitin particles (GCP) driving a mixed Th1/Th17 response (3) and 3) GCP and the TLR4 agonist (GLA) expected to drive a pronounced Th17 response (15). Following three vaccinations two weeks apart, and a rest period of 12 months (**Fig. 2B**), mice vaccinated with the combination of GCP and GLA survived significantly longer compared to mice vaccinated with either Advax3 or GCP (**Fig. 2C**). To uncover the mechanism of the improved survival, we analyzed the development of the T cell responses at serial time points post-vaccination (**Fig. 2B**). At two weeks after the last vaccine boost, both the GCP+GLA and GCP alone groups had increased frequencies and numbers of *Bl*-Eng-2 specific tetramer-positive CD4 T cells in the lungs after challenge that coincided with a reduction in lung CFU compared to unvaccinated controls (**Fig. 2C**). At three months post-boost, after effector T cells had contracted, the GLA group demonstrated increased frequency and number of tetramer-positive memory T cells and reduced lung CFU compared to the GCP vaccinated mice (**Fig. 2C**). In summary, the addition of GLA to the GCP-*Bl-*Eng2 vaccine increased the development of memory CD4^+^ T cells and resistance to infection with *Bd*.

**Fig. 2:**
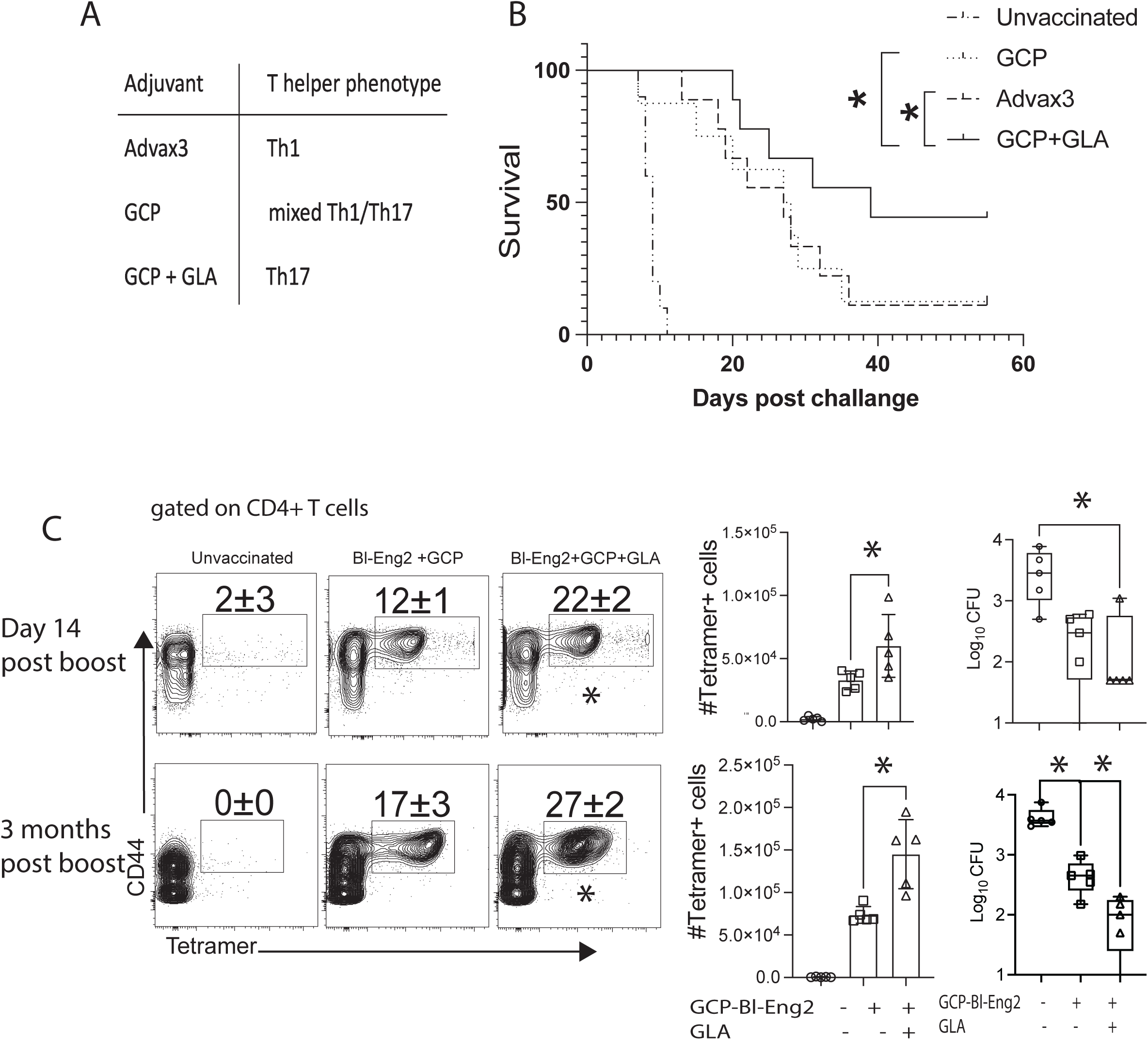
Combination adjuvants augment protective efficacy. Mice were vaccinated with *Bl*-Eng2 and a combination of adjuvants that drive either a polarized Th1, Th17 or mixed Th1/Th17 response (**A**). Vaccination with GCP-*Bl*-Eng2 and GLA provided increased survival compared to the other adjuvant combinations, *p<0.05, Kaplan Meier test (**B**). After two weeks and 3 months post-vaccination, mice were challenged with *Bd* yeast and lung T cells were analyzed for the frequency and number of tetramer positive T cells at day 4 post-infection, *p<0.05, Anova test (**C**).

### Combination adjuvants including GLA induce polyfunctional Th1 cells

To investigate CD4 T cell function, we performed intracellular-cytokine staining of primed T cells after *ex vivo* stimulation with *Bl-*Eng2 peptide and analyzed the T helper phenotypes at the peak of T cell expansion (day 14 post boost) and after contraction (3- and 12-months post boost) (**Fig. 3**). Mice vaccinated with Advax3 exhibited a pronounced Th1 response throughout the time course, whereas the formulation with GCP resulted in a balanced (1:1 ratio) of Th1:Th17 cells (**Fig. 3A**). The addition of GLA yielded a 1:1 ratio at the peak of the T cell expansion (day 14), but a skewed Th1:Th17 ratio of 2:1 by 3 months and 6:1 by 12 months post boost. IFN-γ positive CD4 T cells at 12 months post vaccination also co-expressed TNF and GM-CSF indicating that they were polyfunctional Th1 cells (**Fig. 3B**). The number of Th1 cells that produced one (IFN-γ), two (IFN-γ + TNF or IFN-γ + GM-CSF) or three (IFN-γ + TNF + GM-CSF) cytokines was significantly elevated with the addition of GLA (**Fig. 3C**) compared to the other vaccine groups. From these data, we hypothesized that the addition of GLA either promotes the loss of Th17 cells during the contraction phase or the conversion of Th17 cells into polyfunctional Th1 cells, which are highly effective in protecting against this infection.

**Fig. 3:**
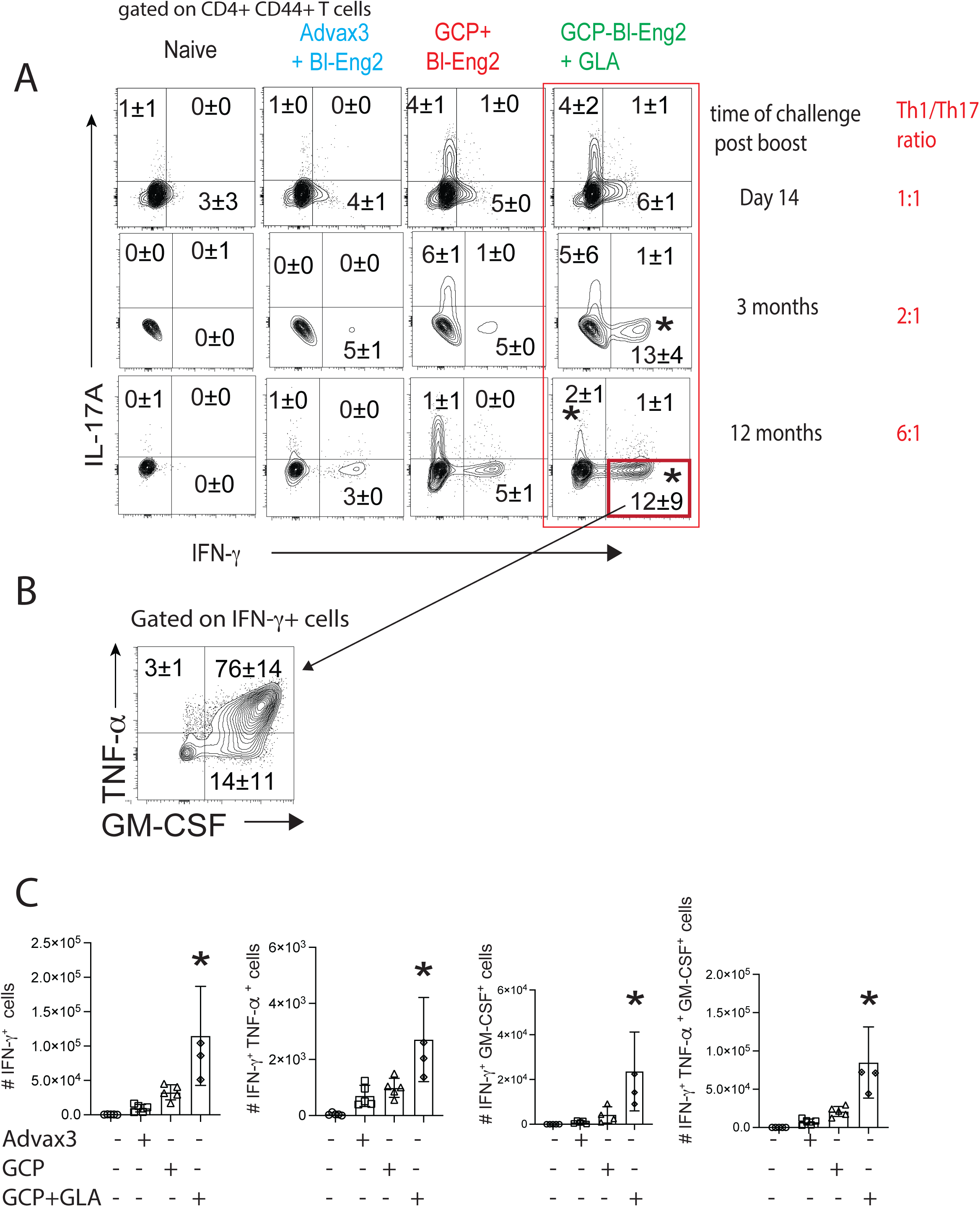
The addition of GLA to GCP-*Bl-*Eng2 induces the generation of polyfunctional Th1 cells after 3- and 12-months post-vaccination. Mice were vaccinated with Advax3+*Bl*-Eng2, GCP-*Bl*-Eng2 or GCP-*Bl*-Eng2+GLA thrice two weeks apart and rested for 14 days or 3 months after the last boost. CD4^+^CD44^+^ T cells were analyzed for intracellular production of IFN-γ and IL-17, *p<0.05 vs. all other groups, Anova test (**A**). IFN-γ+ T cells were analyzed for the production of TNF and GM-CSF 12 months after vaccination. (**B**). The number of single (IFN-γ^+^), double (IFNγ^+^/TNF^+^ and IFNγ^+^/GM-CSF^+^) and triple (IFN-γ^+^/TNF+/GM-CSF^+^) type 1 cytokine producing T cells were enumerated by FACS 12 months after vaccination, *p<0.05, vs. all other groups, Anova test. (**C**).

### Vaccination with *Bl*-Eng2 and GLA drives the differentiation of plastic Th17 cells in IL-17 fate-mapping reporter mice

To test the conversion hypothesis, we used IL-17-eYFP fate mapping mice to let us permanently mark IL-17A^+^ cells to see whether Th17 cells do indeed convert into Th1 cells to sustain vaccine protection. To investigate the conversion of Th17 to Th1 cells, we analyzed the cytokine production of *Bl*-Eng2-specific T cells at the peak of T cell expansion (day 14 post-boost) and after contraction (3 months post-boost). At the peak of the T cell response, the frequencies and numbers of IFN-γ, IL-17 (**Fig. 4A**) and reporter-positive T cells (**Fig. 4B**) were similar in the presence and absence of GLA. However, after contraction at 3 months post-vaccination, the addition of GLA during vaccine administration resulted in increased frequencies and numbers of reporter-positive T cells. In addition, in mice vaccinated with GCP-*Bl*-Eng2 + GLA, the frequency of tetramer^+^ T cells was increased among reporter^+^ T cells (18.1%) compared to total CD4^+^ T cells (5.76%) indicating that *Bl*-Eng2-specific cells are increased among the IL-17 lineage^+^ T cells (**SFig. 1A**).

**Fig. 4:**
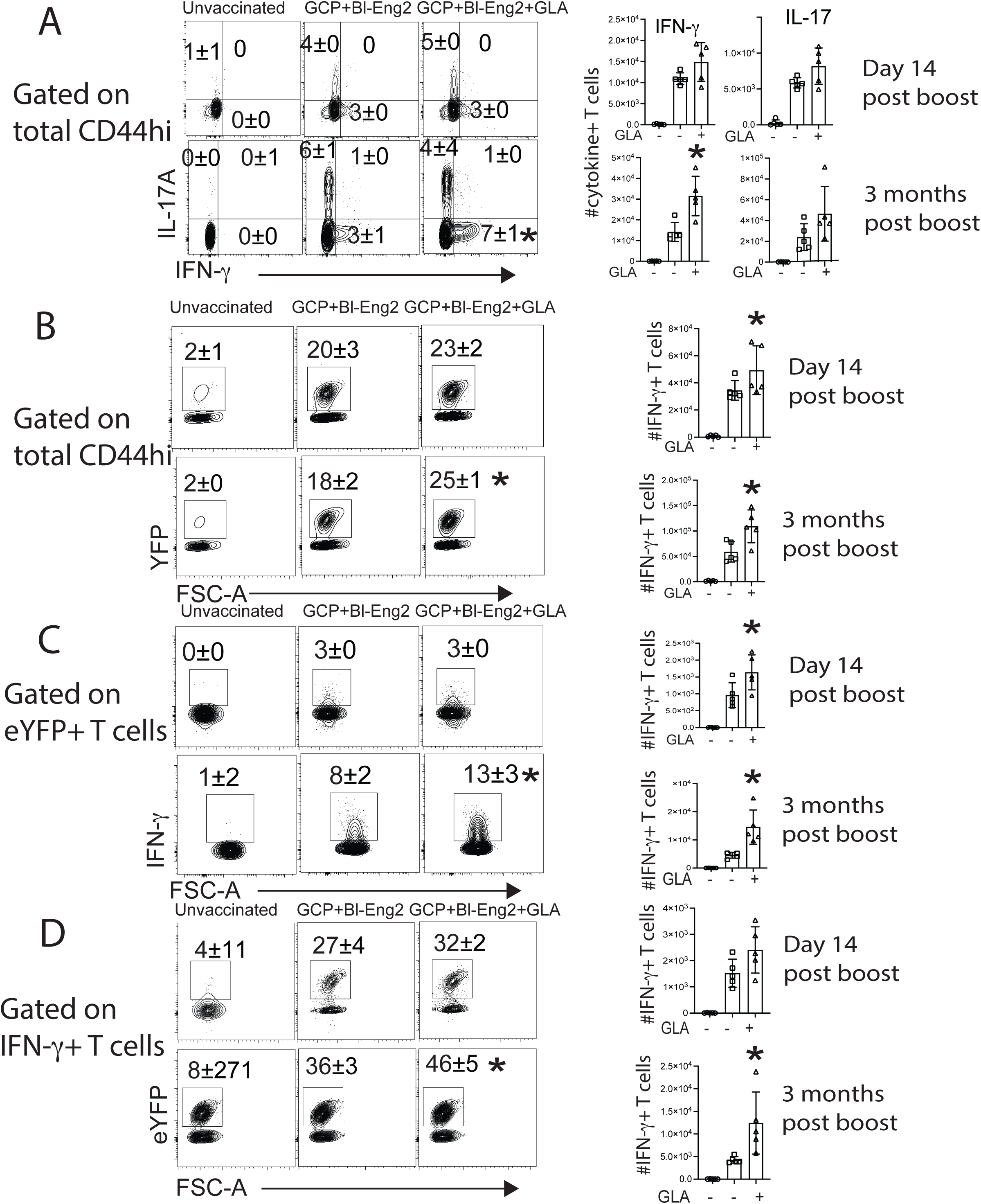
Combination adjuvants with GLA and *Bl-*Eng2 induce plastic Th17 cells in IL-17 fate mapping mice that convert to IFN-γ producing Th1 cells. IL-17 reporter (fate mapping) mice were vaccinated with GCP-Eng2 or GCP-*Bl-*Eng2+GLA and lung T cells analyzed at two weeks and three months post-vaccination. CD4^+^CD44^+^ T cells were analyzed for intracellular IFN-γ and IL-17 production (**A**) and eYFP (**B**). eYFP^+^ T cells were analyzed for intracellular IFN-γ (**C**). IFN-γ^+^ T cells were analyzed for eYFP fluorescence (**D**). *p<0.05, vs. all other groups, Anova test.

To investigate whether Th17 cells convert to Th1 cells, we gated on eYFP positive T cells and analyzed the expression of IFN-γ. The addition of GLA increased the frequencies and numbers of IFN-γ-expressing Th17 cells at 3 months post-vaccination (**Fig. 4C**). In addition, the frequency of IFN-γ and IL-17 expressing T cells was increased among eYFP^+^ T cells compared to eYFP^-^ and total CD4^+^ CD44^+^ T cells in mice vaccinated with GCP-*Bl*-Eng2+GLA (**SFig. 1B**). IFN-γ^+^ T cells emerging from the IL-17 T cell lineage also expressed both GM-CSF and TNF at a high frequency (>80%) indicating that the majority of the reporter-positive cells are polyfunctional (**SFig. 1B**). Similarly, IFN-γ^+^ T cells from the eYFP negative lineage were also largely polyfunctional. To determine the percentage of IFN-γ expressing T cells that stem from the Th17 cell lineage, we gated on IFN-γ-positive T cells and analyzed the expression of eYFP. Almost half of the IFN-γ expressing T cells originated from the Th17 cell lineage in mice that were vaccinated with GLA (**Fig. 4D**). To summarize, these data indicate that GLA promotes the development of plastic Th17 cells that convert into long-lasting, multi-functional, memory Th1 cells.

### Plastic Th17 cells convert to Th1 cells during contraction/memory phase

In mice immunized with GCP+*Bl-Eng2*+GLA, we found above that Th17 cells convert into Th1 cells after contraction of T cells by 3 months after immunization (**Fig. 3**). We also found that *Bl-*Eng2-specific Th17 cells had converted to Th1 cells after *Bd* challenge following 3 months of rest post-immunization (**Fig. 4**). This finding could result from the conversion of the T17 (to Th1) helper phenotype either during the 3 months of rest or the secondary expansion in lung after infection. To resolve this question, we assessed the T helper profiles of Eng2-specific T cells in vaccinated mice immunized both <u>before</u> and <u>after</u> infection. We analyzed T cells from the spleen before infection and lung T cells at day 4 post-infection (**Fig. 5**). We had previously shown that Eng2- specific T cells migrate from the spleen to the lung after challenge and mediate protection (3). While the frequency of tetramer positive T cells was higher in the lungs after challenge than in the spleen before challenge (**Fig. 5A**), the number of Eng-2 specific T cells was much higher in the spleen (**SFig. 2**). The frequencies of eYFP^+^ T cells before and after challenge was similar (**Fig. 5B**) which allowed us to compare the frequencies of the T helper phenotypes and conversion of Th17 (reporter positive) cells into Th1 cells. Among eYFP^+^ T cells, 16% from unchallenged mice and 8.8% from challenged mice expressed IFN-γ (**Fig. 5C**). Among CD4^+^ CD44^+^ T cells 4% from unchallenged mice and 2.6% from challenged mice expressed IFN-γ (**Fig. 5D**). Among IFN-γ positive T cells 38% from unchallenged mice and 24% from challenged mice stemmed from eYFP^+^ T cells (**Fig. 5E**). These results indicate that conversion of plastic Th17 cells to Th1 cells occurs before challenge during the 3 months period of contraction/memory and not during secondary expansion in the lung after infection.

**Fig. 5:**
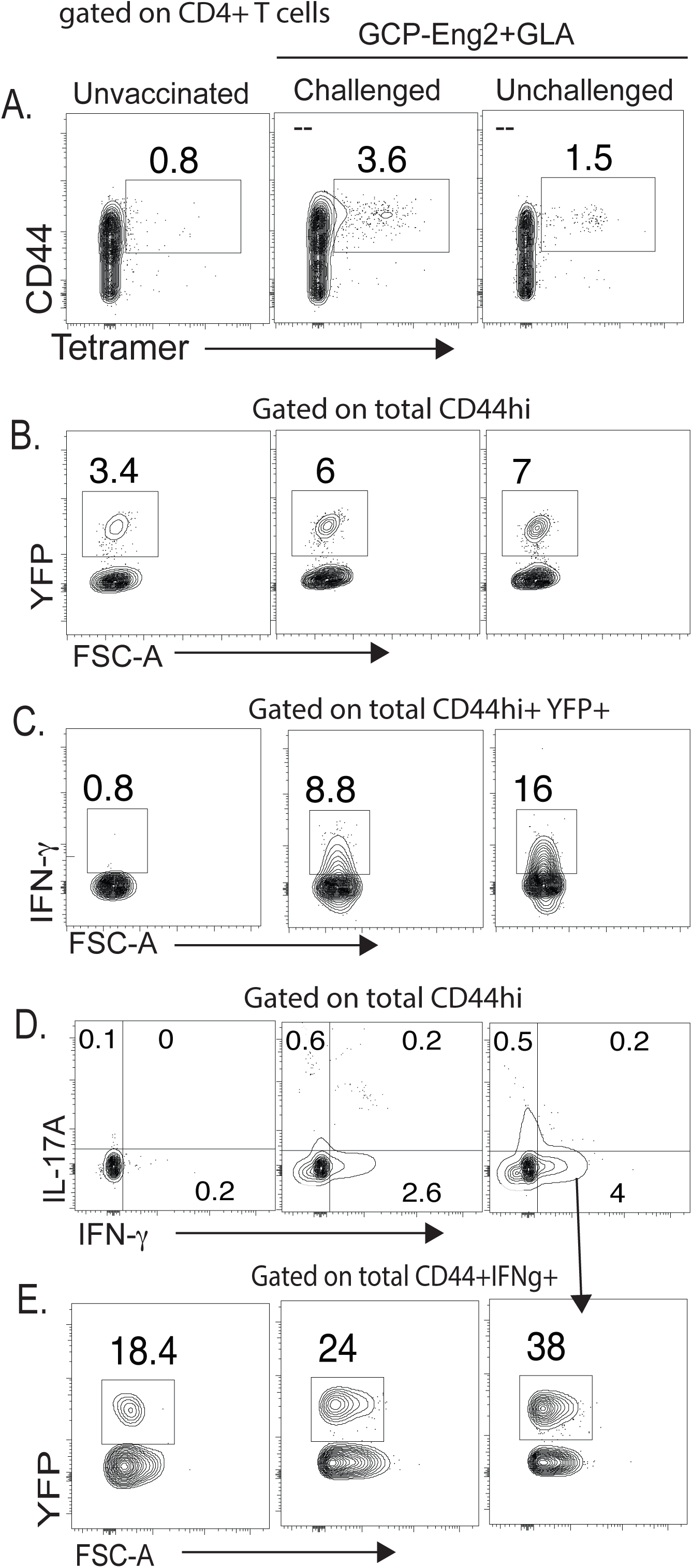
The conversion of plastic Th17 cells to Th1 cells occurs during the 3 months contraction/memory phase. Mice were vaccinated with GCP-Eng2+GLA thrice two weeks apart. At 3 months post-vaccination we harvested splenocytes from unchallenged mice and lung T cells from challenged mice at day 6 post-infection. Dot plots display the frequencies of tetramer^+^ cells (**A**), eYFP^+^ cells (**B**), IFN-γ^+^ T cells among eYFP^+^ T cells (**C**), IFN-γ and IL-17^+^ T cells among CD4^+^ CD44^+^ T cells (**D**) and eYFP reporter^+^ T cells that produced IFN-γ (**E**).

### Adoptive transfer of plastic Th17 cells protects naïve mice against fungal infection

To investigate whether plastic Th17 cells that convert to Th1 cells mediate anti-fungal resistance we employed two experimental approaches: 1) we adoptively transferred eYFP reporter positive T cells from vaccinated mice that have been rested for 3 months post-vaccination into naïve recipient mice and 2) we adoptively transferred eYFP^+^ T cells at the peak expansion (day 14 post-infection) and let them rest in the recipient mice for 3 months to allow conversion to Th1 cells.

Prior to adoptive transfer, we characterized the donor cells from mice that were vaccinated with GCP-*Bl-*Eng2+GLA and rested for 3 months post-vaccination. We challenged a subset of these mice and analyzed the recall response in the lung at day 4 post-infection. Among CD4^+^ T cells 31.5% were tetramer-positive and among CD4^+^ CD44^+^ T cells 40.7% were reporter-positive (**SFig. 3A**). Of the tetramer positive cells, 54.5% were reporter positive and, among the reporter positive cells, 56.4% were tetramer positive. When stimulated with *Bl*-Eng2 peptide, of the cytokine producing reporter-positive T cells, two thirds produced IL-17 and one third produced IFN-γ, whereas reporter negative T cells produced mostly IFN-γ (**SFig. 3B**). Lung CFU in vaccinated donor mice were significantly reduced compared to unvaccinated control mice (**SFig. 3C**). Thus, cells from donor mice were effective in controlling infection prior to transfer.

For adoptive transfer, we sorted CD4^+^, eYFP positive T cells from the spleen of unchallenged donor mice and transferred them into naïve, wild type recipients. The next day we challenged the recipients and analyzed transferred eYFP^+^ cells and tetramer^+^ cells recalled to lung. The frequencies and numbers of eYFP^+^ and tetramer^+^ T cells were enriched in the lungs of recipients compared to controls (**Fig. 6A-C**). This enrichment coincided with reduced lung CFU at day 4 post-infection in the recipients compared to mice that did not receive eYFP^+^ T cells (**Fig. 6D**). In a second cohort of mice, we adoptively transferred eYFP^+^ cells and determined lung CFU at the time that naïve recipients who did not receive cells became moribund. Adoptive transfer of eYFP positive cells likewise reduced lung CFU significantly (**Fig. 6E**). These data indicate that plastic Th17 cells that become multifunctional are protective against fungal infection.

**Fig. 6:**
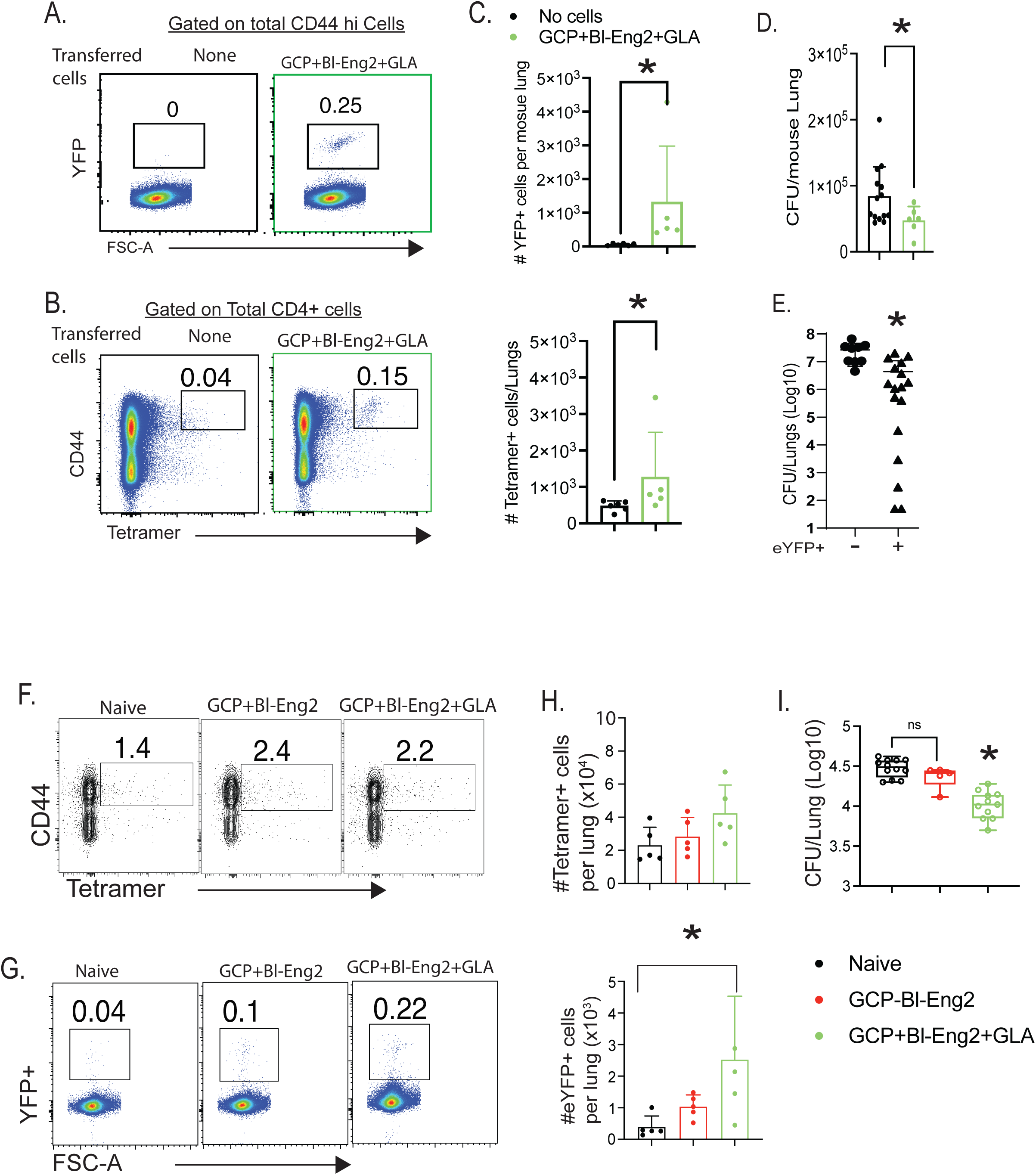
Adoptive transfer of eYFP^+^ reporter cells after vaccination protects recipient mice against fungal infection. Donor mice were vaccinated with GCP-*Bl*-Eng2+GLA thrice. The mice were rested for 3 months post-vaccination allowing memory cells to form within these donor mice. At day 4 post-infection, lung CD4^+^ T cells were enriched from these mice by positive selection, sorted for eYFP^+^ T cells and the cells adoptively transferred. Naïve wild type recipient mice were challenged the next day and lung T cells (**A-C**) and CFU analyzed at day 4 post-infection (**D**), * p<0.05, 2-tailed Mann-Whitney U test. A separate cohort of recipient mice was analyzed for lung CFU when control mice that did not receive T cells became moribund (day 14 post-infection) (**E**). eYFP^+^ T cells were adoptively transferred 14 days post-vaccination. Memory cells were allowed to form in the recipients for 3 months before challenge (**F-I**). The lungs of recipients were harvested 4 days post-infection. Transferred T cells were analyzed by FACS (**F-H**) and lung CFU enumerated (**I**). *p<0.05, vs. all other groups, Anova test.

In the second approach, we transferred cells at the peak of expansion and rested them in recipients. Here, we vaccinated mice with GCP-*Bl*-Eng2 or GCP-*Bl*-Eng2+GLA and adoptively transferred eYFP^+^ T cells at day 14 post-vaccination and let the T cells rest for 3 months to allow conversion to Th1 cells in the recipient mice. The frequencies and numbers of eYFP^+^ and tetramer^+^ T cells were enriched in the lungs on recall in both groups of recipient mice compared to controls (**Fig. 6F-H**). Importantly, the addition of GLA to vaccination increased the number of eYFP^+^ and tetramer^+^ T cells upon recall, which coincided with reduction of lung CFU compared to recipients that either did not receive cells or received them from GCP-*Bl*-Eng2 vaccinated mice (**Fig. 6I**). Thus, the results from both experimental approaches – either resting cells with plasticity and conversion occurring over 3 months in either the donors or the recipients – indicate that plastic Th17 cells adoptively transferred protection to naïve recipient mice.

### Polyfunctional Th1 cells mediate vaccine induced protection

We investigated the relative contributions to vaccine resistance of IL-17 or IFN-γ independently as compared to IFN-γ, TNF and GM-CSF combined. We neutralized these cytokines in mice that were vaccinated with GCP-*Bl-*Eng2+GLA and had been rested for 9 months. Neutralization of either IL-17 and IFN-γ alone increased lung CFU by one or two logs, respectively (**Fig. 7**). However, the combined neutralization of IFN-γ, TNF and GM-CSF together abolished vaccine induced protection entirely and increased lung CFU at least to the level of unvaccinated control mice. These results indicate that IL-17 and IFN-γ partially contribute to vaccine resistance, whereas polyfunctional Th1 cells that produce IFN-γ, TNF and GM-CSF are most effective in rendering this type of resistance.

**Fig. 7:**
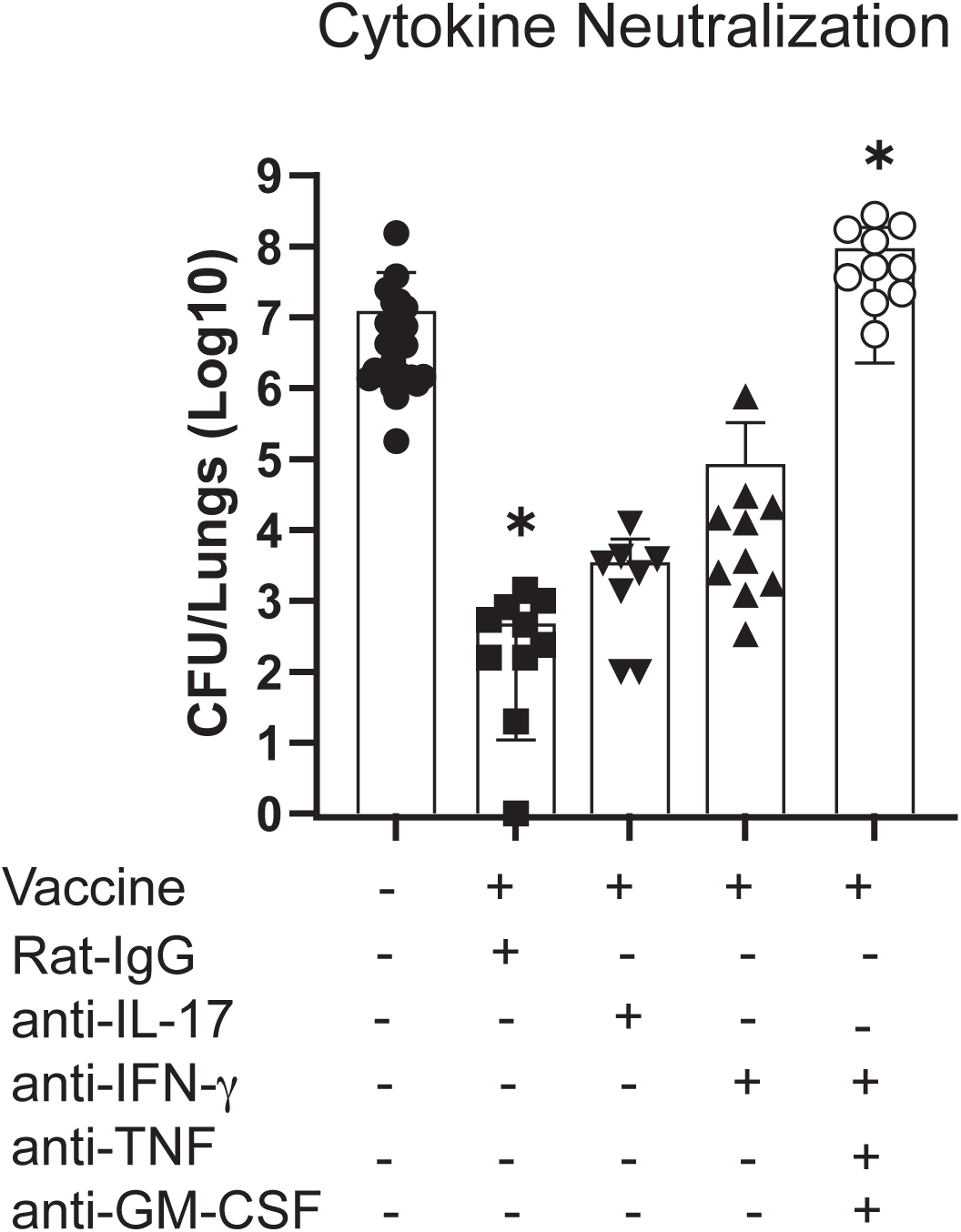
Mice were vaccinated with GCP-*Bl*-Eng2+GLA thrice and rested for 9 months post-vaccination. At the time of challenge and every other day thereafter vaccinated mice were treated with 250 μg of antibodies as indicated. At day 9 post-infection when unvaccinated mice were moribund, lung CFU were plated. * p<0.05 vs. all other groups, Anova test.

## DISCUSSION

The pivotal role of IFN-γ and type 1 cytokines in shaping immunity to fungi and other intracellular microbes makes understanding the role and action of adjuvant combinations that drive Th1 immunity germane for harnessing the immune system for therapeutic benefit. Additionally, multifunctional T helper cells have been linked to better vaccine induced immunity against the intracellular pathogen *Mycobacterium tuberculosis* (*16*). Herein, we exploited a unique combination of the fungal antigen Eng-2 GLA with several adjuvants that included GLA, a ligand for TLR4, to enhance resistance to fungi in a manner that persuades CD4^+^ T cells to become mighty producers of IFN-γ and other type 1 cytokines. The addition of GLA increased protective efficacy against *Bd* by promoting the development of plastic Th17 cells that yielded polyfunctional Th1 memory T cells and accelerated vaccine development. While GLA boosted the numbers of plastic Th17 cells, both Th17 and non-Th17 cell lineages contributed to the development of polyfunctional Th1 cells.

The plasticity of vaccine induced T cells was unexpected in our study. We had previously published that vaccine induced anti-fungal Tc17 (CD8^+^ T cells) were not plastic and were maintained as stable, long-lasting memory cells that resisted conversion into IFN-γ producing cells (Tc1) that protected CD4^+^ T cell deficient hosts against lethal pulmonary fungal infection (17). We observed here that combination adjuvant with GLA in the subunit vaccine formulation induced Th17 (IL-17 producing CD4^+^) cells that converted to Th1 (IFN-γ producing) T cells. Thus, while both vaccine-induced anti-fungal CD4^+^Th17 and CD8^+^ Tc17 cells persist long term – for at least one year - they differ in their ability to convert to IFN-γ producing T cells. In contrast to anti-fungal Tc17 cells, anti-viral Tc17 cells behave differently. For example, vaccination with Influenza nucleoprotein (NP) and the synthetic TLR4 agonist GLA induced long term memory Tc17 cells (unpublished data) that, upon viral re-challenge, converted to protective Tc1 effector cells. Thus, long-term anti-viral immunity is facilitated by the stem-like behavior of Tc17 cells in vaccinated mice, which convert to effector Tc1 cells during infection when Tc1 cells are needed to combat influenza. In contrast to the timing of conversion of anti-viral Tc17 cells to Tc1 cells, *Bl*-Eng2 specific plastic Th17 cells converted to Th1 cells during the contraction and memory phases during rest after vaccination rather than during the secondary expansion phase following re-infection as seen for anti-viral Tc17 cells.

Th17 cells reportedly display context-dependent plasticity, as they are capable of acquiring functional characteristics of Th1 cells (18). The plasticity of this subset is associated with higher *in vivo* survival and self-renewal capacity and less senescence than Th1 polarized cells, which have less plasticity and more phenotypic stability (18). This late plasticity may contribute to their protection against microbes, while also being necessary for antitumor activity of Th17 cells in adoptive cell transfer therapy models (18); however, this behavior also is thought to play a role in the development of autoimmunity (18).

The stability or plasticity of Th subsets is governed by the epigenetic regulation of the key transcription factors and cytokines determining the polarization status. Whether a particular gene is poised for expression is determined by the chromatin structure, histone and DNA methylation states (19). Trimethylation of histone ^3^H on lysine 4 (H3K4me3) is considered permissive for gene expression, whereas trimethylation of lysine 27 on histone ^3^H (H3K27me3) is a marker of gene silencing. In some cases, both states can be found in a gene locus, thus making it susceptible for either expression or negative regulation. Gene promoters for *tbx21* in Th1 cells and *rorc* in Th17 cells displayed a permissive methylation state (H3K4me3) associated with full expression of these master regulators in each lineage. However, gene promoters for *tbx21* in Th17 cells had a bivalent status characterized by H3K4me3/H3K27me3 dual positivity, thus substantiating the relative instability and propensity of these cells to acquire type 1 features in the presence of IL-12 (20). In vaccination induced EAE almost all myelin-specific CD4^+^ T cells infiltrating CNS were of the Th17 origin but ceased secretion of IL-17 and switched to producing IFN-γ (21). This *in vivo* conversion was critically dependent on IL-23. In contrast, in acute cutaneous candidiasis, responding Th17 cells remained firmly committed to IL-17 production, possibly because of low local levels of IL-23.

The finding that T17 cells converted to multifunctional T cells during the contraction phase after vaccination was unexpected. During the contraction phase, 90–95% of activated effector T cells die, resolving inflammation and preventing immunopathology. This phase follows pathogen or antigen clearance, leaving behind a small number of long-lived memory cells. The conversion of Th17 cells to Th1 cells is generally believed to occur during active inflammation, especially in the setting of autoimmune disease where the cells mediate pathogenesis of disease (22). Such conversion generally requires inflammatory cytokines such as IL-12, IL-23 or IL-1β. We did not investigate the presence or participation of these cytokines in mediating conversion of plastic Th17 cells into Th1 in our study. However, we previously reported that p35^-/-^ and IL-12rβ2^-/-^ mice acquired resistance when vaccinated with a live attenuated strain of *B. dermatitidis,* whereas p40^-/-^ failed to do so (23). These results are compatible with the idea that IL-23 is required for the acquisition of vaccine immunity and conversion of Th17 into Th1 cells. We surmise the combination of adjuvants together (GLA+GCP+Eng2) provide a depot of ongoing, low-level inflammation that persists at the site of vaccination, enabling conversion in the setting of required inflammatory cytokines.

## SUPPLEMENTAL FIGURES

**SFig. 1 for main figure 4: Polyfunctional T cells in IL-17 reporter mice vaccinated with GCP-*Bl*-Eng2+GLA.** Mice were vaccinated with GCP-*Bl*-Eng2+GLA twice and challenged 3 months post-vaccination. The frequencies of tetramer^+^ T cells were determined from total CD4^+^ T cells and CD4^+^ CD44^+^ eYFP^+^ T cells (**A**). The frequencies of cytokine (GM-CSF, TNF and IFN-γ) producing T cells were determined from CD4^+^ CD44^+^ T cells and the frequencies of GM-CSF and TNF producing T cells from IFN-γ^+^ T cells (**B**). *p<0.05, vs. all other groups, Anova test.

**SFig. 2 for main figure 5: Number of tetramer^+^, eYFP^+^ and cytokine producing T cells in unchallenged and challenged reporter mice**. eYFP reporter mice were vaccinated with GCP-Eng2+GLA thrice two weeks apart and rested for 3 months before challenge. Lung T cells were analyzed in challenged reporter mice 6 days post-infection and in splenocytes from unchallenged reporter mice. The number of tetramer^+^ T cells (**A**), eYFP^+^ T cells (**B**), eYFP^+^ IFN-γ^+^ T cells (**C**), IL-17^+^ T cells (**D**), IFN-γ^+^ T cells (**E**) and ex-Th17^+^Th1 cells (**F**) were enumerated. *p<0.05, vs. all other groups, Anova test.

**SFig. 3 for main figure 6:** Properties of donor T cells. Donor T cells were analyzed before adoptive transfer and lung CFU of the donors was enumerated 4 days post-infection. Donor mice were vaccinated with GCP-*Bl*-Eng2+GLA twice and rested for 3 months. Lung T cells were stained for tetramer (**A**) and cytokines (**B**) and analyzed by FACS. Lung CFU were plated from the same mice (**C**). *p<0.05, 2-tailed, Mann-Whitney U test.

## Supporting information

SFig. 1 for main Fig. 4

SFig. 2 for main Fig. 5

SFig. 3 for main Fig. 6

## Acknowledgement

The work was supported by NIH grants BAA-NIAID-DAIT-AI201800007, Contract Number: 75N93019C00064, R24AI192252, R01AI93553 (MW), R01AI040996 (BK/MW), R01 AI168370 (BK), U01 AI124299 (BK), R37 AI035681 (BK), and by the American Heart Association Postdoctoral Fellowship #835129 (LSD). Flow samples were processed at the University of Wisconsin Carbone Cancer Center (UWCCC) Flow Core Facility on a BD LSR Fortessa that was purchased with the NIH shared instrumentation grant 1S100OD018202-01 and University of Wisconsin Carbone Cancer Center Support grant P30 CA014520.

