## Supplementary figures and images for "Combination adjuvants drive long lived plastic Th17 cells that convert to multi-functional Th1 cells and protect mice against fungal infection"

### SFig. 1 for main Fig. 4

#### Supplementary figure 1 main figure 4

##### A. Surface

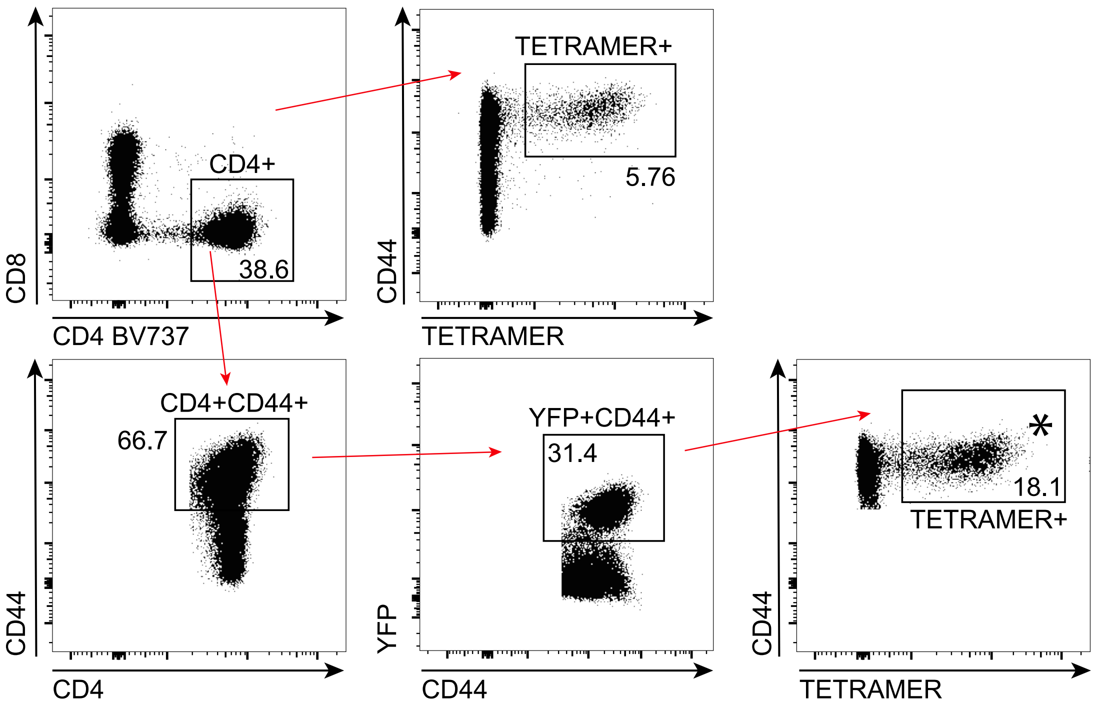


##### B. ICS

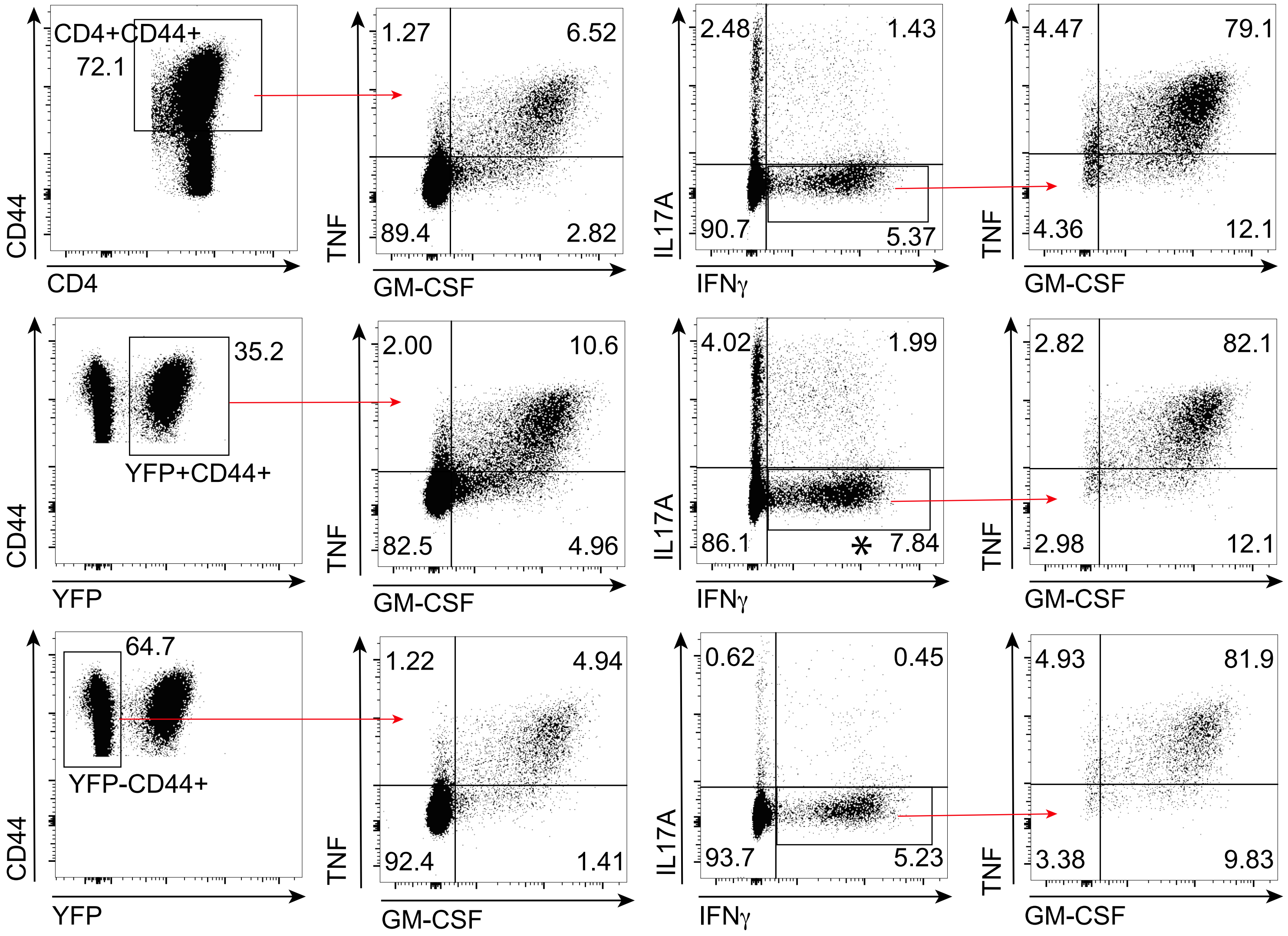

### SFig. 2 for main Fig. 5

#### SFig. 2 for main figure 5

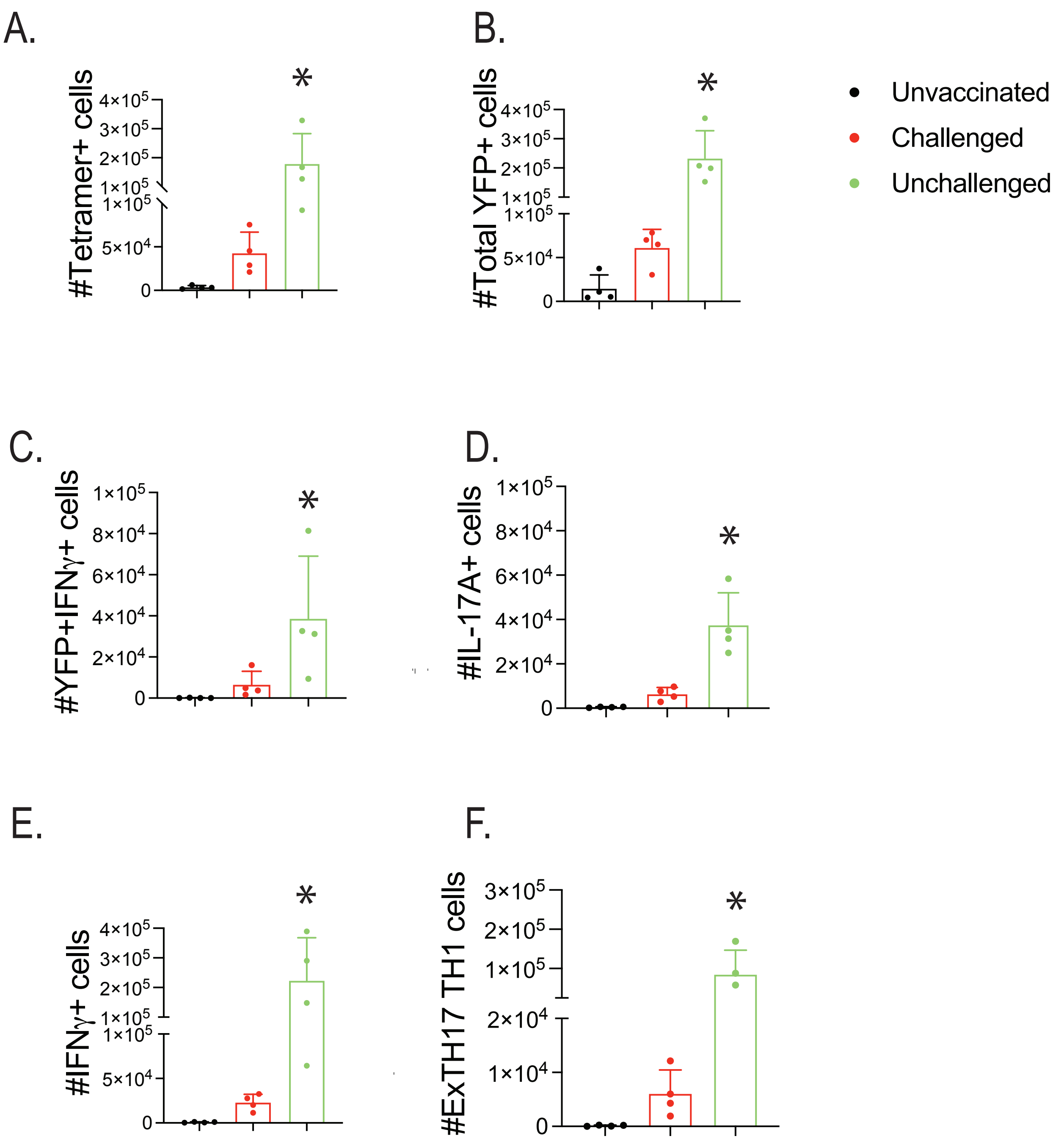

### SFig. 3 for main Fig. 6

#### SFig. 3 for main figure 6

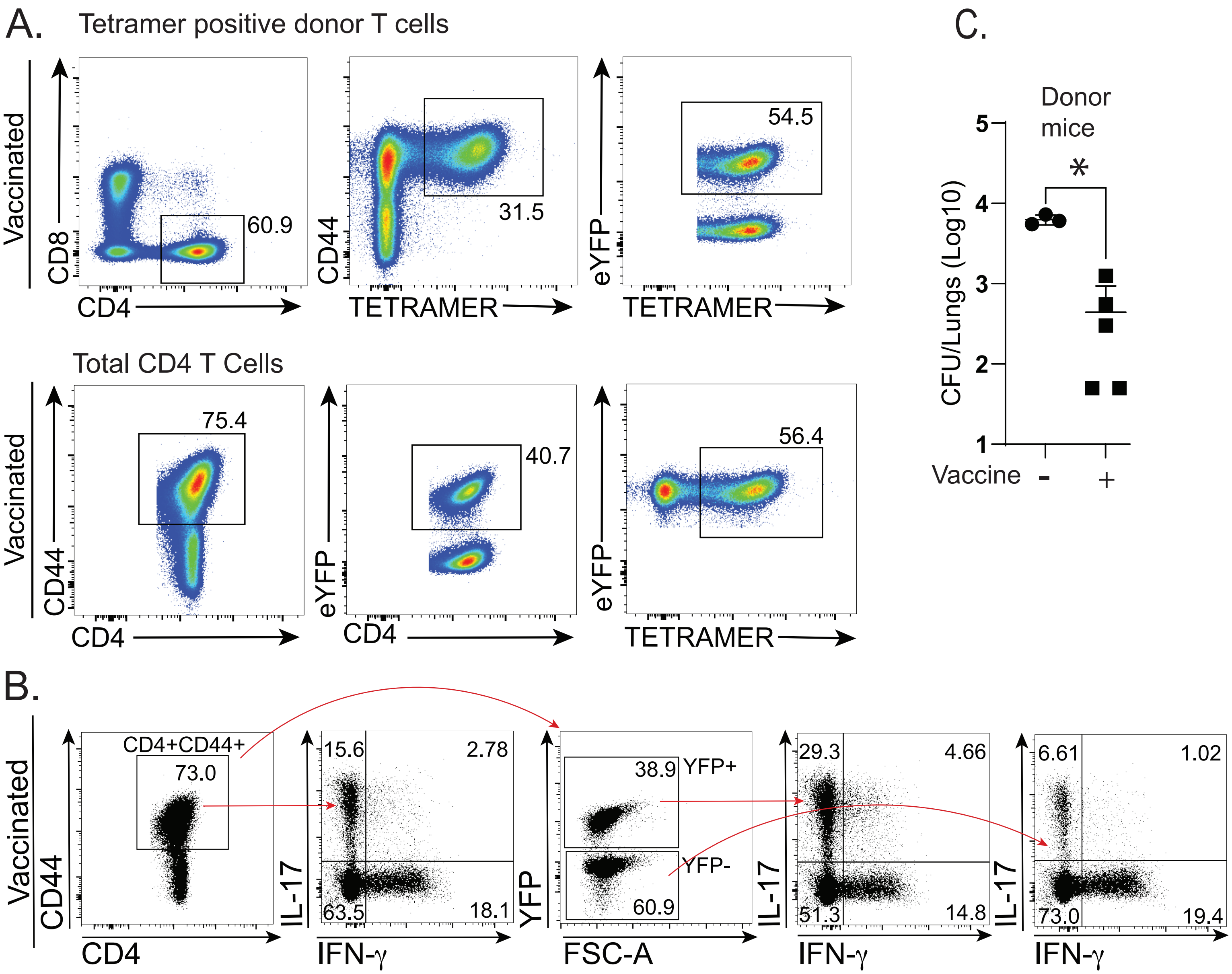
